# A Multiscale Coarse-grained Model of the SARS-CoV-2 Virion

**DOI:** 10.1101/2020.10.02.323915

**Authors:** Alvin Yu, Alexander J. Pak, Peng He, Viviana Monje-Galvan, Lorenzo Casalino, Zied Gaieb, Abigail C. Dommer, Rommie E. Amaro, Gregory A. Voth

**Affiliations:** Department of Chemistry, Chicago Center for Theoretical Chemistry, Institute for Biophysical Dynamics, and James Franck Institute, The University of Chicago, Chicago, IL 60637; Department of Chemistry and Biochemistry, University of California San Diego, La Jolla, CA 92093

**Keywords:** SARS-CoV-2, virion, coarse-graining, molecular dynamics simulations, multiscale

## Abstract

The severe acute respiratory syndrome coronavirus 2 (SARS-CoV-2) is the causative agent of the COVID-19 pandemic. Computer simulations of complete viral particles can provide theoretical insights into large-scale viral processes including assembly, budding, egress, entry, and fusion. Detailed atomistic simulations, however, are constrained to shorter timescales and require billion-atom simulations for these processes. Here, we report the current status and on-going development of a largely “bottom-up” coarse-grained (CG) model of the SARS-CoV-2 virion. Structural data from a combination of cryo-electron microscopy (cryo-EM), x-ray crystallography, and computational predictions were used to build molecular models of structural SARS-CoV-2 proteins, which were then assembled into a complete virion model. We describe how CG molecular interactions can be derived from all-atom simulations, how viral behavior difficult to capture in atomistic simulations can be incorporated into the CG models, and how the CG models can be iteratively improved as new data becomes publicly available. Our initial CG model and the detailed methods presented are intended to serve as a resource for researchers working on COVID-19 who are interested in performing multiscale simulations of the SARS-CoV-2 virion.

**Significance Statement:** This study reports the construction of a molecular model for the SARS-CoV-2 virion and details our multiscale approach towards model refinement. The resulting model and methods can be applied to and enable the simulation of SARS-CoV-2 virions.

## Introduction

The onset of the global COVID-19 pandemic has brought intense investigation into the molecular components of SARS-CoV-2 encoded by the virus’s 30 kilobase (kb) genome. Structural biology efforts using cryo-electron microscopy (cryo-EM) and x-ray crystallographic techniques are currently reporting new structures of viral proteins every week (1–12), and computational structure prediction efforts are targeting unresolved sections of the genome using a variety of protein folding algorithms. Computational and experimental studies are underway to find new molecular therapeutics that can inhibit viral activity or further elucidate the mechanisms of action of SARS-CoV-2 proteins (13–16). The computer simulation of large-scale SARS-CoV-2 processes, such as virion assembly, budding, entry, and fusion, will remain intrinsically challenging to investigate using all-atom (AA) molecular dynamics (MD), owing to the computational cost of meaningfully simulating the hundred of millions to billions of atoms involved.

A holistic model of the SARS-CoV-2 virion can provide insight into the mechanisms of large-scale viral processes and the collective behavior of macromolecules involved in viral replication and infectivity. SARS-CoV-2 virions contain four main structural proteins: the spike (S), membrane (M), nucleocapsid (N), and envelope (E) proteins (17). S proteins are glycosylated trimers that mediate fusion and entry, in part by attaching enclosed fusion peptide sequences into the membranes of host cells (18). M proteins appear as dimeric complexes embedded within the virion envelope, and are believed to anchor ribonucleoprotein complexes to the envelope (19, 20). N proteins associate with and organize RNA into ribonucleoprotein structures found in the interior of virions (21, 22). Lastly, E proteins are thought to form pentameric ion channels that are found at the lipid bilayers of virion membranes and contribute to viral budding (23).

In this paper, we construct a largely “bottom-up” coarse-grained (CG) model of the SARS-CoV-2 virion from the currently available structural and atomistic simulation data on SARS-CoV-2 proteins. In general, this model serves as a resource for researchers working on COVID-19, and as a platform to incorporate computational and experimental data. This model also enables the multiscale study of SARS-CoV-2 processes for the development of treatment and prevention strategies against COVID-19. Atomistic trajectory and experimental structural data deposited in the NSF Molecular Sciences Software Institute (MolSSI) will be incorporated, as they become publicly available (24). In this work, we detail several of our CG methods used to iteratively develop a CG model for the full SARS-CoV-2 virion, in which molecular interactions between CG particles are derived using a combination of phenomenological, experimental, and atomistic simulation approaches.

## Theory and Methodology

### Building models from structural data

We first constructed atomic models of the structural proteins of the SARS-CoV-2 virion, (Figure 1). AA models of the open and closed state of the S protein were built based on the cryo-EM structures of the spike ectodomain (PDB ID: 6VYB, 6VXX) (5), respectively, and atomic models of the N protein were constructed based on the x-ray crystallographic structure of the nucleocapsid NTD (PDB ID: 6M3M) (25). Glycosylation sites were modeled using Glycan Reader & Modeler in CHARMM-GUI (26) and the site-specific glycoprofile derived from mass spectrometry and cryo-EM analysis (27, 28). Homology models for the S protein stalk, including the HR2 and TM domains were assembled as α-helical trimeric bundles using MODELLER (29) on the basis of secondary structure assignments in JPred4 (30). Homology models for the SARS-CoV-2 N protein CTD were created from the x-ray crystallographic structure of the SARS-CoV N protein CTD (PDB ID: 2CJR) (31). Missing amino acid backbones in loop regions were built in MODELLER, and side chain rotamers were built using SCWRL4 (32). For the M protein dimer and the pentameric E ion channel, we used atomic models derived in the recent Critical Assessment of Structure Prediction (CASP) Commons competition (33, 34).

**Figure 1.**
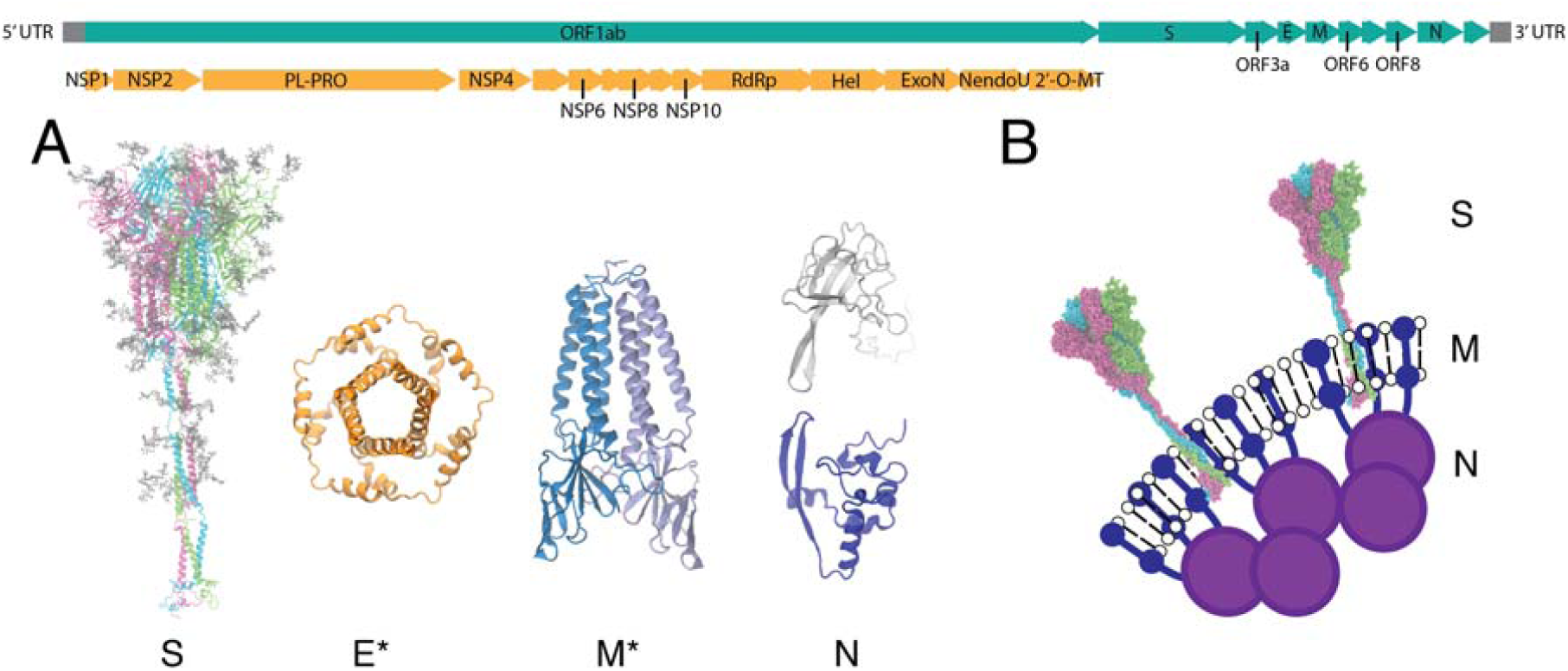
Viral proteins of SARS-CoV-2. The genome of SARS-CoV-2 is shown in the top panel. Nonstructural proteins (NSPs) encoded in the open reading frame (ORF) 1ab are colored in orange, and the full genome is in teal. **(A)** All-atom models of the structural proteins of SARS-CoV-2 consisting of the S, E, M and N proteins. Asterisks indicate homology modeled protein structures for E and M (34). **(B)** Schematic of the virion surface from cryo-EM images of the virion, adapted from Ref. (19).

AA protein models (see discussion below) were subsequently simulated and coarse-grained to generate the CG models (Figure 2 and see sections below). A previously developed CG model for lipids was used, consisting of three CG beads per lipid and distinct bead types for lipid head groups and hydrophobic tails (35). A single component CG lipid bilayer was generated in a spherical configuration and equilibrated using CG molecular dynamics simulations under constant NVT conditions in LAMMPS (36). We note that in the future more complex CG lipid models (37) can be added. Transmembrane segments of component membrane proteins were visually identified and assigned based on secondary structure motifs. Individual lipids on the outer leaflet of the spherical bilayer were randomly selected and used as initial positions for embedding spike, membrane, and envelope proteins. For each initial position, the center-of-mass of the transmembrane domain was aligned with the center of the lipid bilayer, and the principal axis of the protein was aligned with the vector normal to the lipid bilayer. Transmembrane regions were then substituted for the overlapping CG lipids to embed the proteins. The procedure was iterated until a spike, membrane, and envelope protein density on the virion surface was achieved that was approximately consistent with current available experimental estimates of ∼25, 1000, and 20 per virion, respectively, from cryo-EM and biochemical data (38–40). The diameter of the membrane envelope is approximately 100 nm and 120 nm including the S proteins on the virion surface. As higher-resolution experimental data are released, the overall structure of this model can be refined.

**Figure 2.**
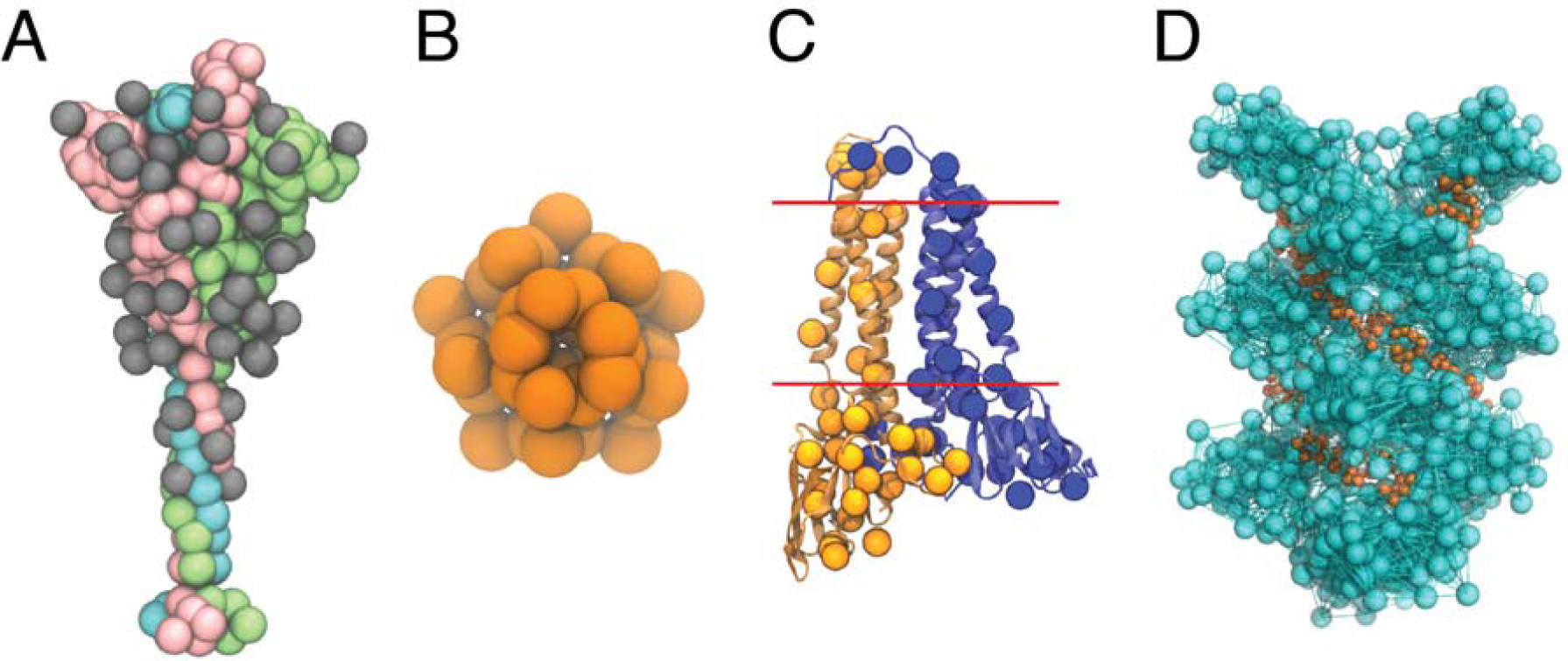
CG models of the SARS-CoV-2 structural proteins. **(A)** The CG model of the S protein trimer in the open state. The protein monomers are depicted as pink, green, and cyan beads, respectively; the monomer in pink has an exposed receptor binding domain. Each of the 22(x3) N-linked glycans are depicted as grey beads. **(B)** The CG model of the pentameric E protein is depicted as orange beads. **(C)** The CG M dimer model is depicted as yellow and blue spheres, overlaid on top of the AA model of the M dimer. Each monomer has 36 CG sites, and the red lines indicate the approximate positions of the transmembrane region. **(D)** The CG model of the N protein CTD helix in complex with viral RNA. The N protein helix and bonds derived from the hENM are depicted in cyan, while the RNA is depicted as orange beads.

### AA MD simulations of the S protein

Two glycosylated models of the open and closed spike were inserted into a symmetric 225 Å × 225 Å lipid bilayer mimicking the composition of the endoplasmic reticulum-Golgi intermediate compartment (41, 42). The lipid patch was built using CHARMM-GUI. The complete protein-membrane system was solvated with using the TIP3P water model (43), and neutralized with chloride and sodium ions to maintain a 150 mM concentration. Each system contained ∼1.7 million atoms. Minimization and equilibration was performed using the CHARMM36 forcefield (44, 45) and NAMD 2.14 (46). Production runs were performed in the NPT ensemble using a Langevin thermostat at 310K and Nosé-Hoover Langevin barostat at 1 atm. All production runs used a 2-fs timestep and the SHAKE algorithm. Multiple replicas of AA MD simulations of the open (3x) and closed (3x) systems were performed on NSF Frontera at the Texas Advanced Computing Center (TACC), achieving an aggregate sampling of 3.0 and 1.8 µs, respectively.

### CG model of the S protein

The CG model of the glycosylated S protein (Figure 2 A) was parameterized from the AA MD simulations described below (47). Reference statistics used conformations sampled equally from both open and closed states with AA trajectories spanning the 3.0 and 1.8 µs, respectively. First, the protein was mapped to CG beads using essential dynamics coarse-graining (EDCG) (48). We used 60 and 50 CG beads for the S_1_ and S_2_ domains, respectively, and the 22 N-linked glycans were each mapped to a single bead. Intra-protein interactions were represented as a hetero-elastic network model (hENM) with bond energies *k* (*r* − *r*_0_)^2^ where *k* is the spring constant of a particular CG bond and *r*_0_ is the equilibrium bond length. These parameters were optimized using the hENM method (49). Inter-protein interactions within the S-trimers were composed of excluded volume, attractive, and screened electrostatic terms. For excluded volume interactions, a phenomenological soft cosine potential, 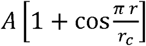, was used, where *A* = 25 kcal/mol and *r*_*c*_ is the onset for excluded volume. Attractive, non-bonded interactions between inter-protein contacts were modeled as the sum of two Gaussian potentials, 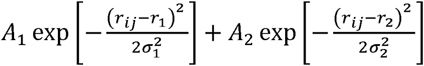, where *r*_1_ and *σ*_1_ are the mean and standard deviation determined by a fit to the pair correlation between CG sites *i* and *j* through least-squares regression. The constants. *A*_1_ and *A*_2_, were optimized using relative-entropy minimization (REM). Screened electrostatics were modeled using Yukawa potentials, 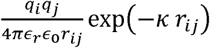, where *q*_*i*_ is the aggregate charge of CG site *i, k* = 1.274 nm^−1^ is the inverse Debye length for 0.15 M NaCl, and *ϵ*_*r*_ is the effective dielectric constant of the protein environment, approximated as 17.5 (50).

### CG models of the M and E proteins

AA simulations of the M protein dimer were performed using homology models and a membrane model based on the endoplasmic reticulum (ER). The membrane model included PC:PE:PI:PS:Chol lipids (0.45:0.10:0.23:0.10:0.12 mole fraction) as an initial approximation to the ER-Golgi intermediate compartment (ERGIC). The protein-membrane systems were solvated and neutralized in a similar fashion as described previously. The simulations were equilibrated for 400 ns prior to a 4 µs production run on Anton2. All simulations were run in the constant NPT ensemble at 310K and 1 atm using the CHARMM36m forcefield. A CG model containing approximately 5 residues per CG bead was mapped from the reference statistics of the AA MD simulations using the EDCG (Figure 2C), and hENM approaches. A CG model for the E protein was developed by linearly mapping the amino acid sequence to particles at a resolution of 1 CG bead per 5 amino acids (Figure 2B).

### CG model of the N protein

Several studies suggest that the C-terminal domain (CTD) of the N protein assembles into a helix that contains two RNA binding grooves (21, 51). Based on these studies, we constructed atomic and CG models of the viral ribonucleoprotein complex (vRNP) by iterating between CG and AA simulations. We first constructed an atomic model of the N protein CTD helix with two RNA binding grooves by stacking 3 copies of the CTD octamer structure (PDB: 2CJR), which is composed of 4 CTD dimers and homology modeled from the X-ray crystallographic structure of the SARS-CoV N protein CTD (31). The CTD helix was simulated in the CHARMM36m force field for 400 ns. We then constructed the CTD helix model using EDCG combined with hENM followed by manually placing CG RNA beads into the groove of the helix (Figure 2D). The positions of the CG beads were used as restraints to build an atomic model of the vRNP complex. Finally, the vRNP model was relaxed and simulated in the CHARMM36m force field for 400 ns. It is important to note that recent cryo-EM studies have found granule-like densities within the virion for the vRNP complex (22). Structural detail into how CTD oligomers (including the previously proposed helical model) and RNA fit into these densities will likely require higher-resolution images.

### Deriving CG molecular interactions from AA simulations

Several computational approaches have been developed to build or refine CG models using data from AA or fine-grained simulations. Our approach to coarse-graining the SARS-CoV-2 virion is to couple several CG methods in a hierarchical fashion. CG sites or “beads” are mapped from atomic structures using EDCG, a method designed to preserve the principal modes of motion sampled during atomic-level simulations (48). In EDCG, a given CG mapping operator, 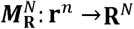, that relates the configurations of the atomistic trajectory (**r**^*n*^) to that of the CG model (**R**^*N*^), is variationally optimized using simulated annealing. Typically, the mapping is constructed so that contiguous segments of a protein’s primary amino acid sequence are mapped to distinct CG sites. For a fixed number of CG beads, *N*, the sets of atoms that are mapped to CG sites are adjusted to minimize the target residual:

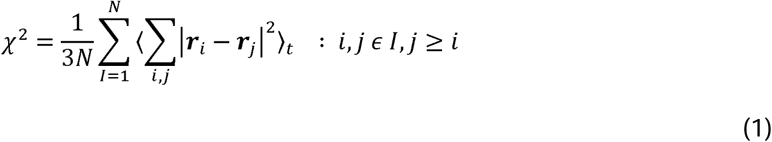

where, *I* = 1, …, *N* is the CG site index, the brackets, ⟨·⟩_*t*_, denote a time-averaged quantity, the sum over *i, j* is a sum over all unique pairs in the set of atoms belonging to the CG site, *I*, and ***r***_*i*_ = ***x***_***i***_ − ⟨***x***_***i***_⟩_*t*_ is the displacement of atom ⟨***x***_***i***_⟩_*t*_ from the atom’s mean position, Note that the residual is small when the displacements, ***r***_*i*_ and ***r***_*j*_, are similar, i.e. the motions of atoms in the same CG site are correlated. A new map is constructed and either accepted or rejected according to a Metropolis-Hastings criterion (i.e., accepted if 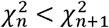; otherwise accepted or rejected such that the new map has probability 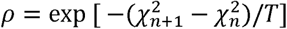, where *n* is the number of iterations for simulated annealing, and is the coupling to a fictitious temperature, that is gradually lowered during optimization).

After defining the AA ↔ CG map, intramolecular interactions within a single polypeptide chain are treated using elastic network models (ENMs) to capture protein flexibility. In the hENM method (49), effective harmonic bonds are assigned to all pairs of particles in the CG model within a tunable distance cutoff that all initially have the same force constant, *k*_*ij*_, between particle *i*, and *j* to construct the bonded topology of the CG model. The harmonic force constants are optimized by first computing the normal modes of this model. In other words, solving the eigenvalue problem,

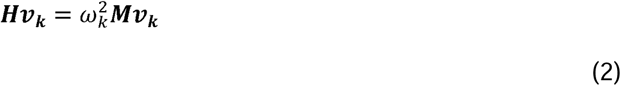

where, *H* is the Hessian: 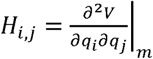, ***M*** is the diagonal matrix for the masses of the particles, and *ω*_*k*_ frequency for the mode of motion. Note that this is the solution to the equation of motion

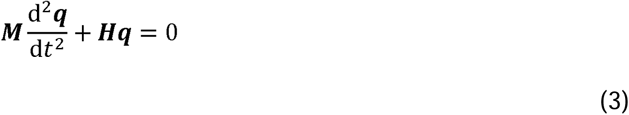

where ***q*** is the generalized coordinate, and that for *N* classically, interacting particles near the potential energy minimum, ***q***_*m*_:

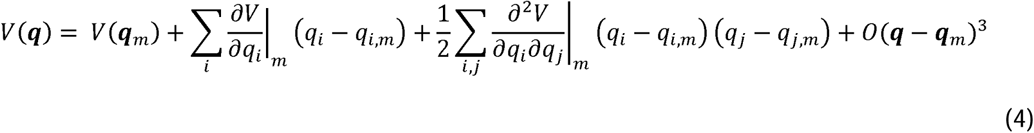

*V*(***q***_*m*_) is constant, and 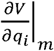 is zero. Using the normal modes, mean-squared fluctuations 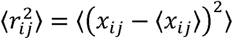 for each *i, j* pair can be computed by rescaling the amplitudes according to an equipartition of energy that reflects the temperature of the atomistic data. Harmonic force constants for each bond in the CG ENM are then iteratively adjusted so that fluctuations in the CG model, match that of the atomistic data, i.e.,

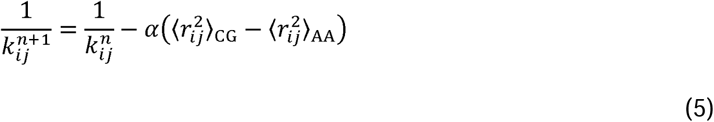

where *n* is the number of iterations and *α* is a parameter that controls the magnitude of the adjustment for each iteration.

For the intermolecular interactions between proteins, non-bonded CG interactions are determined either using force matching (aka multiscale coarse-graining, or MS-CG) (52, 53) or REM approaches (54, 55). In MS-CG, the CG potential is constructed from a linearly independent basis set

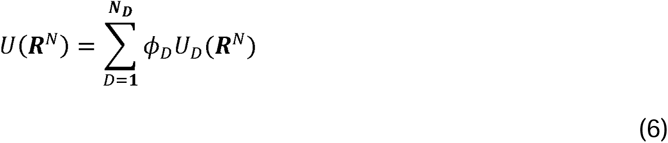

where the functional forms for the basis potentials, *U*_*D*_ (e.g., B-splines, Lennard-Jones, etc), and the number of them, *N*_*D*_, are determined by the user. The coefficients, {*ϕ*_*D*_}, are variationally optimized such that the following residual is minimized:

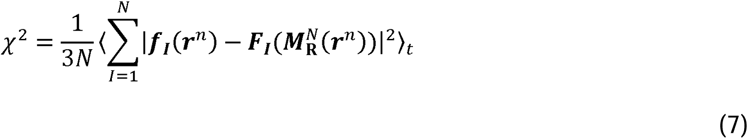

where ***F***_***I***_(***R***^*n*^) = − **∇***U* (***R***^*n*^) is the CG force and ***f***_***I***_(***r***^*n*^) is the atomistic force on the CG site *I*. Similarly, in the REM approach, the objective function for minimization is the Kullback-Liebler divergence, which provides a metric for the differences between the atomistic and CG probability distributions

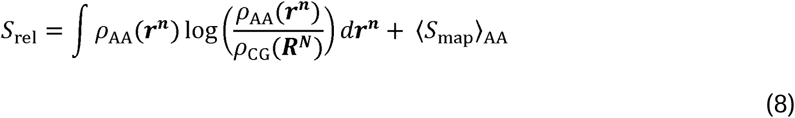

where 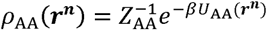 and 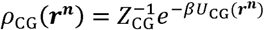 in the canonical ensemble, and *Z* is the configurational partition function. Furthermore, the relative entropy can be expressed as a difference between the potential energy and free energy of the atomistic and CG ensembles:

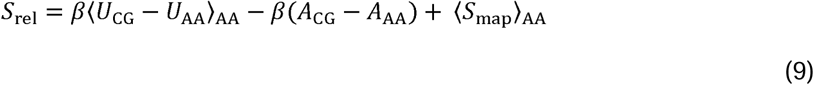

Where *A* = −*k*_B_*T* log *Z*. Minimization of the relative entropy is performed using iterative Newton-Raphson techniques. It is important to note, however, that the quality and fidelity of such CG models is determined by the molecular behavior sampled in the underlying AA simulations.

### Incorporating new behavior in CG simulations

Macromolecular complexes, such as virions, undergo a wide range of behavior including physical and chemical transitions that will be difficult to capture through AA simulations alone, as well as experimental techniques. This is especially true for processes that involve large conformational changes that are not sampled effectively in AA simulations, whether due to the long timescales required, free energy barriers, or inherent limitations of the simulation force field. For instance, the S protein of SARS-CoV-2 exhibits two proteolytic cleavage sites (at the S1/S2 and S2’ locations) as well as binding to the host cell receptor, angiotensin-converting enzyme 2 (ACE-2). Cleavage and binding events trigger dramatic conformational changes in the spike that result in the insertion of the fusion peptide into the host cell membrane. High-resolution structural studies of the S-ACE-2 complex have made protein binding simulations amenable to enhanced sampling techniques at the AA level (4, 56). The proteolytic cleavage of the spike and large-scale conformational shift towards fusion peptide insertion, however, are more difficult to sample in atomistic simulations. To address these issues, one can use CG molecular simulation techniques that allow CG particles to adaptively switch discrete “states” and interactions, such as “Ultra-Coarse-Graining” (UCG) (57–59). In the limit of infrequent internal state switches, UCG implements microscopically reversible state changes are coupled to a Metropolis Hastings like criterion:

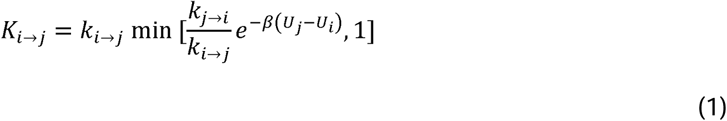

where *K*_*i*→*j*_ is the instantaneous switching rate from state *i* to *j, U*_*j*_ − *U*_*i*_ is the CG effective potential energy difference between state *j* and *i*, and the rates, *K*_*i*→*j*_ and *K*_*j*→*i*_ are model parameters either treated as input or calculated from atomistic simulations. This approach is similar to hybrid kinetic Monte-Carlo and MD methods, but with a spatial kinematic component, and it can be used to examine the transitions of the spike (its “states”) that lead to the fusion of SARS-CoV-2 with host cells.

Experiments can probe longer timescales than are available from AA MD simulations. In recent cryo-EM images of SARS-CoV-2 particles, the S1 domain of the S protein was found to transiently open and close in order to bind the ACE-2 receptor (3, 5), which are subtle conformational changes that are difficult to sample in atomistic simulations. For these conformational changes – in the case that they cannot be treated as discrete state switches – plastic network models (60) or multi-configuration coarse graining (MCCG) methods (61) can be used to construct a CG model that continuously transitions from one state to the next. For plastic network models, two known experimental configurations of the protein are used to build a multi-basin ENM that represents deviations away from each of the individual conformational minima. A phenomenological interaction Hamiltonian is constructed that couples and mixes the ENMs between the two structural endpoints. In MCCG, the primary difference is that the coupling terms in the Hamiltonian are constructed from a two-state mixing approach, derived on the basis of a mapped potential of mean force which is explicitly computed from AA simulation data along collective variables that distinguish between the two (or more) conformational states at a CG level.

### Phenomenological CG models

An alternative (and sometimes necessary “top-down”) route to deriving CG models is to construct a model Hamiltonian and then analyze the model’s resulting behavior in the context of the assumed interactions. Typically, parameterization of such models is designed to fit or reproduce particular observables measured in experimental data and perhaps particular sets of AA simulation data. These can be performed based on variational optimization of some system-specific functional that depends on the experimental observable. Model Hamiltonian approaches have the advantage that physical intuition is clearer, but are not systematic, since each new problem requires a different treatment for the set of interactions involved. Furthermore, these approaches often require orthogonal experiments to validate the underlying model. Such coarse-graining methods are, however, especially useful in cases where atomistic simulation and/or experimental data is difficult or infeasible to obtain on the system or if the bottom-up methods described above are not expected to yield converged results for the CG effective potential, owing to limited atomistic sampling.

### Early Results

Here we present results from the first CG simulations of the SARS-CoV-2 virion (Figure 3). It should be noted that these are early results and we can thus expect additional simulations to become available from this model as more experimental data and AA simulations become available for the various virion components. In addition, the overall CG methodology and modeling of the virion will continue to evolve, and are works in progress.

**Figure 3.**
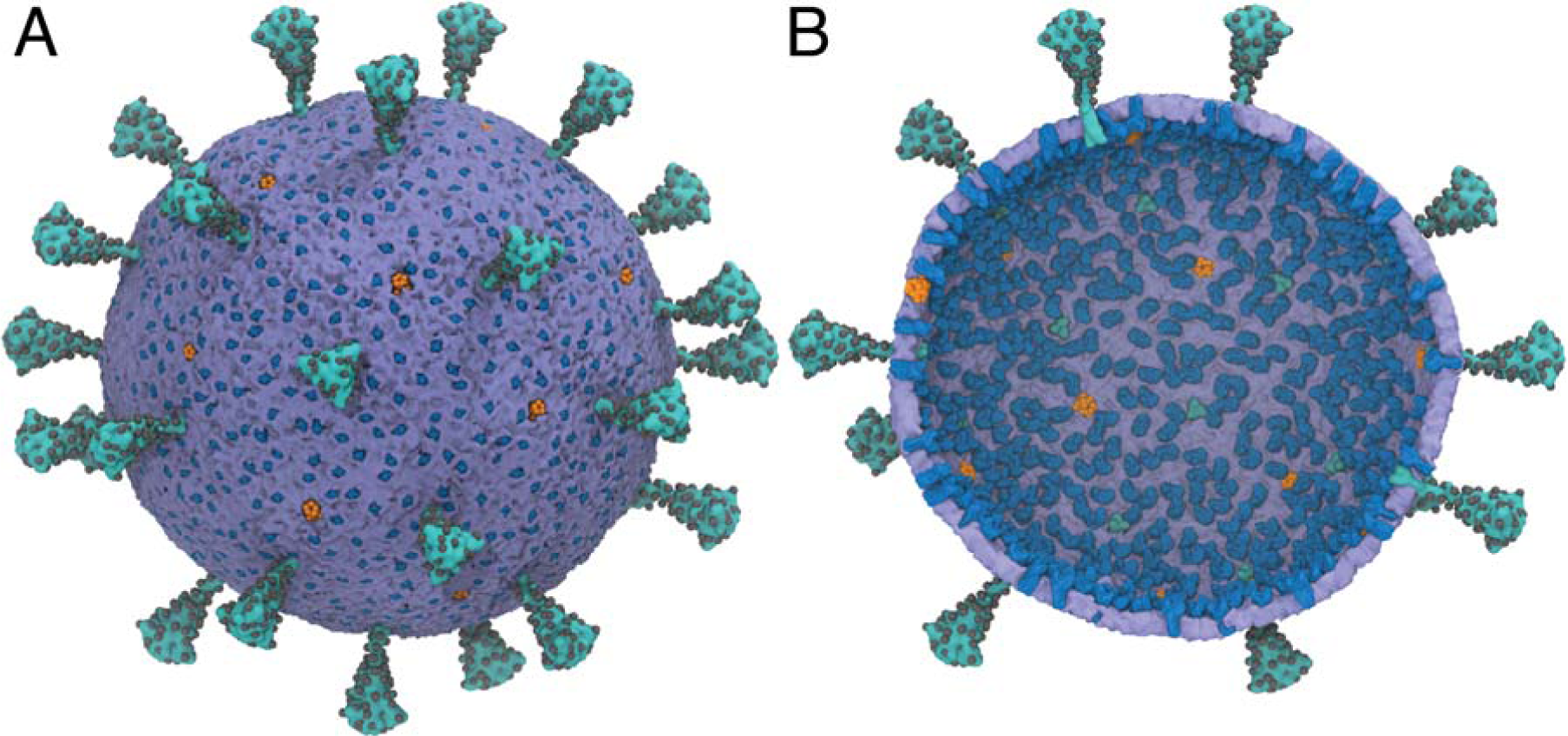
A multiscale model of the SARS-CoV-2 virion. **(A)** Exterior view of the SARS-CoV-2 virion. **(B)** Interior view of the SARS-CoV-2 virion. Spike (S) protein trimers are depicted in teal with the glycosylation sites represented as black spheres. Membrane (M) protein dimers are in blue, with pentameric envelope (E) ion channels in orange. The density of S,M, and E proteins was chosen to be consistent experiments (38–40). N proteins are not shown. The diameter of the membrane envelope is approximately 100 nm and 120 nm including the S proteins on the virion surface.

A CGMD simulation was performed on the complete CG virion model using LAMMPS for 10 × 10^6^ CG timesteps (see Supplementary Movie 1). The system was energy minimized using conjugate gradient descent. A temperature of 300K was maintained with a Langevin thermostat, with a damping constant (*t*_*damp*_ = 10 ps) and 100 fs timestep. Statistics were collected every 100 CG timesteps. Several radial distributions functions (RDFs) or pair correlations between CG particles were computed for the MD trajectories of individual S proteins and compared to the mapped AA reference statistics from which the models were derived (Figure 4 A). In general, the CG model captured the positions and peaks in the pair correlation functions; however, error in the fine structure of the peaks was also present, indicating that refinement involving the addition of more expressive basis CG potentials (e.g., splines) may be necessary.

**Figure 4:**
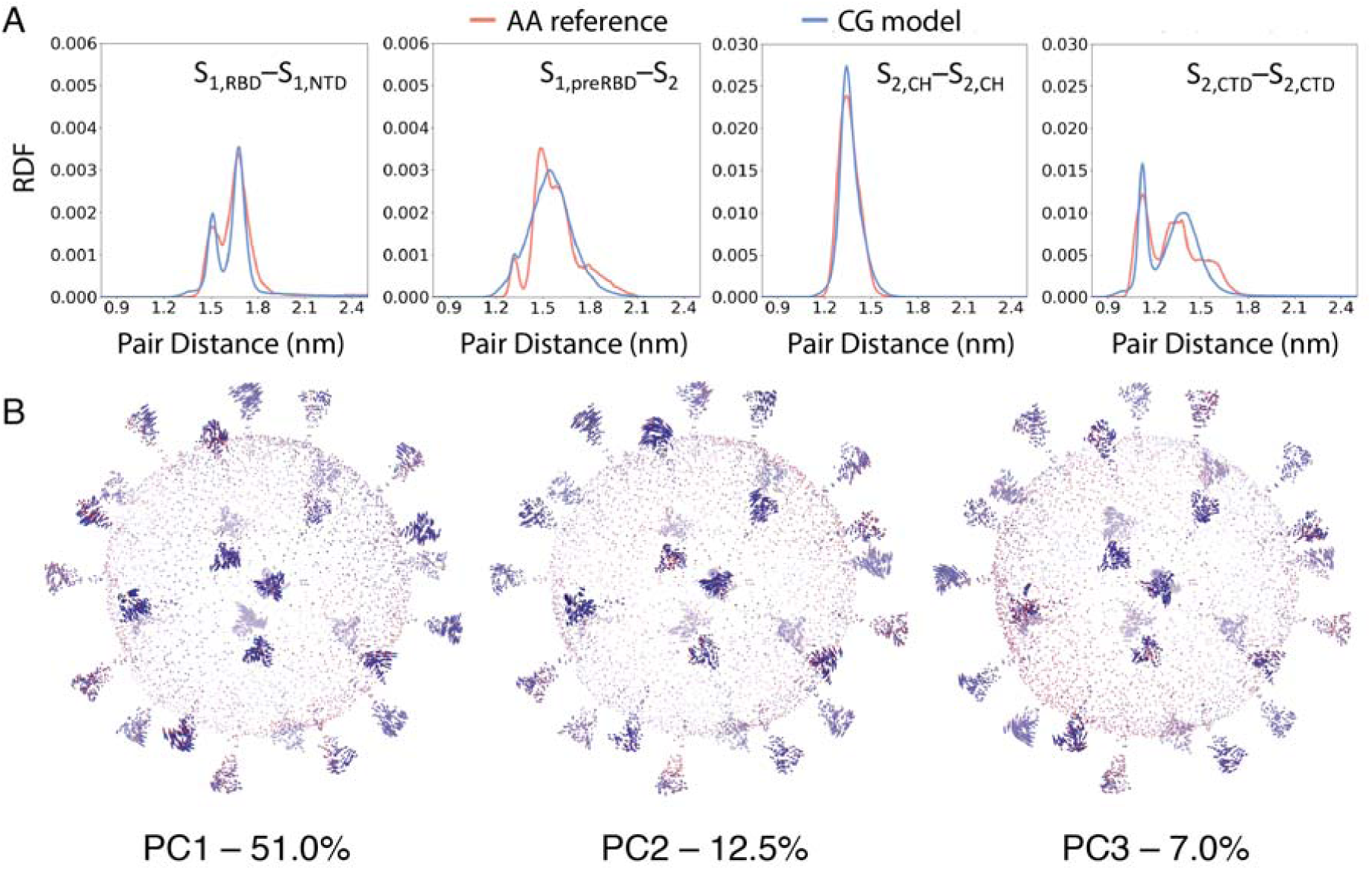
Analysis of the CGMD simulations of the SARS-CoV-2 virion. **(A)** Radial distribution functions (RDFs) showing the comparison between mapped all-atom reference statistics and the CG spike model during the MD simulations. The measured RDFs are for CG particles that were mapped from the following all-atom residues of (i) S_1,RBD_ [S459–D467] and S_1,NTD_ [W104–L118], (ii) S_1,preRBD_ [E309–R319] and S_2_ [A852–L861], (iii) S_2,CH_ [A1015–K1028], and (iv) S_2,CTD_ [Y1215–V1228]. **(B)** Principal modes of motion of the SARS-CoV-2 virion computed from the CGMD simulation (see also Supplementary Movie 2–4). The first principal component (PC1) accounts for 51% of the total variation observed during simulation, whereas the second (PC2) and third (PC3) account for 12.5% and 7%, respectively.

We performed principal component analysis (PCA) on a subset of the CG particles to examine collective modes of motion of the virion (Figure 4B). The Cartesian coordinates of 1 particle for every 15 CG lipids, 1 for every M and E protein, and 1 for every 3 S particles were extracted from the trajectory data and used to compute the covariance matrix, 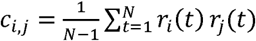, where *r*_*i*_(*t*) is the mean-free position vector, *r*_*i*_(*t*) = *x*_*i*_ − ⟨*x*_*i*_⟩_*t*_ of particle *i*. The highest variance eigenmode, PC1, (see Supplementary Movie 2) corresponds to splaying motions in the S1/S2 domain of the S protein, and accounts for 51% of the total variance seen during the simulations. Similarly, PC2 (see Supplementary Movie 3) accounts for 12.5% of the variance and corresponds to rocking motions of the S1/S2 domain, while PC3 (see Supplementary Movie 4) accounts for 7.0% of the total variance and corresponds to twisting of the S1/S2 during CGMD. In general, there was a high degree of variance in the S protein, and these correlated modes of motion are likely informative of how the virion collectively utilizes spike proteins to explore and detect receptors. Longer CG simulations with more expressive CG models will likely be required to uncover additional modes of motion in the virion, including modes that involve the structural M, N, and E proteins.

### Conclusions and Outlook

This work provides an initial CG molecular model of the SARS-CoV-2 virion and details a bottom-up CG approach capable of further refining the model as new atomistic and experimental data becomes available. Currently, the lipid envelope is described using a particle-based phenomenological model with a soft tunable bending modulus well suited for large-scale membrane deformations, whereas the M and E proteins are modeled as rigid bodies. Intra-spike interactions were developed using relative entropy minimization approaches on the basis of extensive, microsecond AA simulations of the spike protein. The N protein is modeled on the basis of AA simulations of helical oligomers in complex with RNA. Cross-interactions between the lipids and structural proteins are modeled using attractive Gaussian potentials between the hydrophobic lipid tails and the transmembrane domains of membrane proteins. This virion model will be iteratively refined and improved as structural, biochemical, and AA trajectory data are publicly released. The construction of an integrated CG model from individual atomistic simulations will also be benefited by new developments in systematic methods for ensuring consistency between CG models developed from the reference statistics of those simulations. In particular, methods that variationally optimize in a “divide and conquer” fashion on the basis of joint statistics will likely improve model fidelity. Nonetheless, despite these noted challenges, we find that the behavior of SARS-CoV-2 structural proteins is coupled in the virion.

Coarse-grained simulations of viral processes have helped elucidate a wide range of mechanisms in viruses. For example in HIV, CG simulations contributed to the understanding of the self-assembly of the capsid (62) and innate immune sensor recognition and block of viral activity (63), as well as its inhibition by drug molecules (64). Atomistic simulations of ligand binding have also revealed a variety of unexpected, drug-targetable protein-ligand interaction sites (65–71). It is likely that molecular probes into the processes the involving a holistic model of the SARS-CoV-2 virion can reveal new therapeutic strategies that exploit viral mechanisms involving large-scale behavior.

## Author Contributions

A.Y., A.J.P., P.H., V.M.G. and G.A.V. designed research. A.Y. performed modeling and analysis on the virion. A.Y. and A.J.P. performed modeling and analysis on the S protein. A.Y. and P.H. performed modeling and analysis on the N protein. A.Y. and V.M.G. performed modeling and analysis on the M protein. L.C., Z. G., A.C.D, and R.E.A. contributed all-atom simulation data on the S protein. A.Y., A.J.P., P.H., V.M.G., L.C., Z.G., A.C.D., R.E.A., and G.A.V. wrote the paper.

## Acknowledgements

This work was supported in part by the National Science Foundation through NSF RAPID grant CHE-2029092 (A.J.P, P.H., and G.A.V), in part by the National Institute of General Medical Sciences of the National Institutes of Health through grant R01 GM063796 (V.M.G. and G.A.V.), and in part by NIH GM132826, NSF RAPID MCB-2032054, an award from the RCSA Research Corp., and a UC San Diego Moore’s Cancer Center 2020 SARS-COV-2 seed grant (L.C., Z. G., A.C.D, and R.E.A.). Alvin Yu gratefully acknowledges support from the National Institute of Allergy and Infectious Diseases of the National Institutes of Health under grant F32 AI150208. Alexander J. Pak gratefully acknowledges support from the National Institute of Allergy and Infectious Diseases of the National Institutes of Health under grant F32 AI150477. Computational resources were provided by the Research Computing Center at the University of Chicago, Frontera at the Texas Advanced Computer Center funded by the National Science Foundation grant (OAC-1818253), and the Pittsburgh Super Computing Center through the Anton 2 machine under Grant R01GM116961 from the National Institutes of Health, and the specific allocation PSCA17046P. The Anton 2 machine at PSC was generously made available by D.E. Shaw Research.

## Data Availability

The CG virion model is available at: https://doi.datacite.org/dois/10.34974%2Fq8ya-wh69 and https://github.com/alvinyu33/sars-cov-2-public. The model will be periodically updated with new versions as data is added and the model refined,

