## Supplemental Text for "A Multiscale Coarse-grained Model of the SARS-CoV-2 Virion"

Department of Chemistry

The University of Chicago

5735 S. Ellis Ave, SCL 123

Chicago, IL 60637

**Supplementary Movie 1:** CGMD simulation of the SARS-CoV-2 Virion for 1 x 10^6^ CG timesteps.

**Supplementary Movie 2:** Mode of motion of the virion along principal component 1

**Supplementary Movie 3:** Mode of motion of the virion along principal component 2

**Supplementary Movie 4:** Mode of motion of the virion along principal component 3
